# Conditionally rare taxa contribute but do not account for prokaryotic community changes in soils

**DOI:** 10.1101/165001

**Authors:** Rachel Kaminsky, Sergio E. Morales

## Abstract

Conditionally rare taxa (CRT) are thought to greatly impact microbial community turnover across many environments, but little is known about their role in soils. Here, we investigate the contribution of CRT to whole community variation over space and time in a series of geographically distinct soils dedicated to three agricultural practices of differing intensities and sampled over a full seasonal cycle. We demonstrate that soil CRT do not account for observed total community changes, but that these rare taxa can be modified by spatiotemporal filters.

---

Rare species are pervasive in microbial consortia (Curtis et al., 2002). Known as the “rare biosphere,” these members are being increasingly recognized for their importance in microbial communities (Pedrós - Alió, 2007, Lynch and Neufeld, 2015). There are several potential causes of rarity including transience, competition, niche breadth and predation of abundant taxa (Jousset et al., 2016). It has been suggested that the rare biosphere is a dormant “seed bank” wherein members become abundant after predation events (Lennon and Jones, 2011). However, rare microbes can also be active (Campbell et al., 2011, Hugoni et al., 2013, Kurm et al., 2017), signifying several potential ecological roles for the rare biosphere. Recent work identified conditionally rare taxa (CRT) — microbes that are rare at certain points in time / space and “bloom” to abundance at other points — as major contributors to community dynamics in several environments (Shade et al., 2014). This confirms that some rare microbes are transient and are potentially responsible for changes in community structure. In soils, studies have found that rare microbes bloom after disturbances (Aanderud et al., 2015, Fuentes et al., 2016). Despite these important findings, the literature is still developing with regard to the soil rare biosphere. Given the importance of soil microbial communities to mediating ecosystem processes, understanding the contribution of rare soil microbes is of great importance.

We investigated the contribution of conditionally rare prokaryotes using agricultural soils as a model system (see Supplementary Methods). We sampled soils from 24 sites under three agricultural practices in New Zealand. Sampling occurred during three time points over a year to capture the most divergent seasonal stages (summer to winter and return to summer) to assess community changes over space and time. We tested two hypotheses: (H_1_) CRT contribute disproportionately to whole community structure and (H_2_) recruitment from the rare biosphere is linked to spatiotemporal filtering.

Time responsive CRT constituted 4-6% of OTUs in each site while space responsive CRT represented 1% of the total community (*b* value, > 0.9, relative abundance > 0.01%, Figure S1-3). To assess the contribution of CRT to whole community structure, we constructed three Bray-Curtis distance matrices from OTU tables for each site: (1) only OTUs identified as CRT, (2) whole communities including CRT and (3) whole communities excluding CRT. Mantel tests between distance matrices for CRT and whole communities including CRT are insignificant (Figure 1A). The correlation between space sensitive CRT and the whole community including CRT is significant, but weak (R^2^ = 0.02, *P* = 0.002). Mantel tests between communities excluding CRT and whole communities including CRT have R^2^ values close to 1 (Table S1). This indicates that CRT do not contribute significantly to community variation, which does not align with previous findings, though it should be noted that these studies used moving windows analysis (Shade et al., 2014) and measurements of activity (Aanderud et al., 2015), which were not employed here. It is possible that soil CRT have a limited role in soil functions, remaining mostly dormant and only blooming to abundance in extreme cases, as it is estimated that a substantial portion of microbes are inactive (Lennon and Jones, 2011). On the other hand, CRT might be K-selected, investing in few members and therefore exhibiting a life strategy that isn’t reflected in whole community dynamics. This may be favorable given the heterogeneity of soil, wherein CRT may not overtake dominant taxa, but perform key functions that are costly for those taxa. For example, *Desulfosporosinus* is estimated to perform the majority of sulfate-reduction in peatlands despite its relatively minor contribution to community variance (Pester et al., 2010).

**Figure 1.**
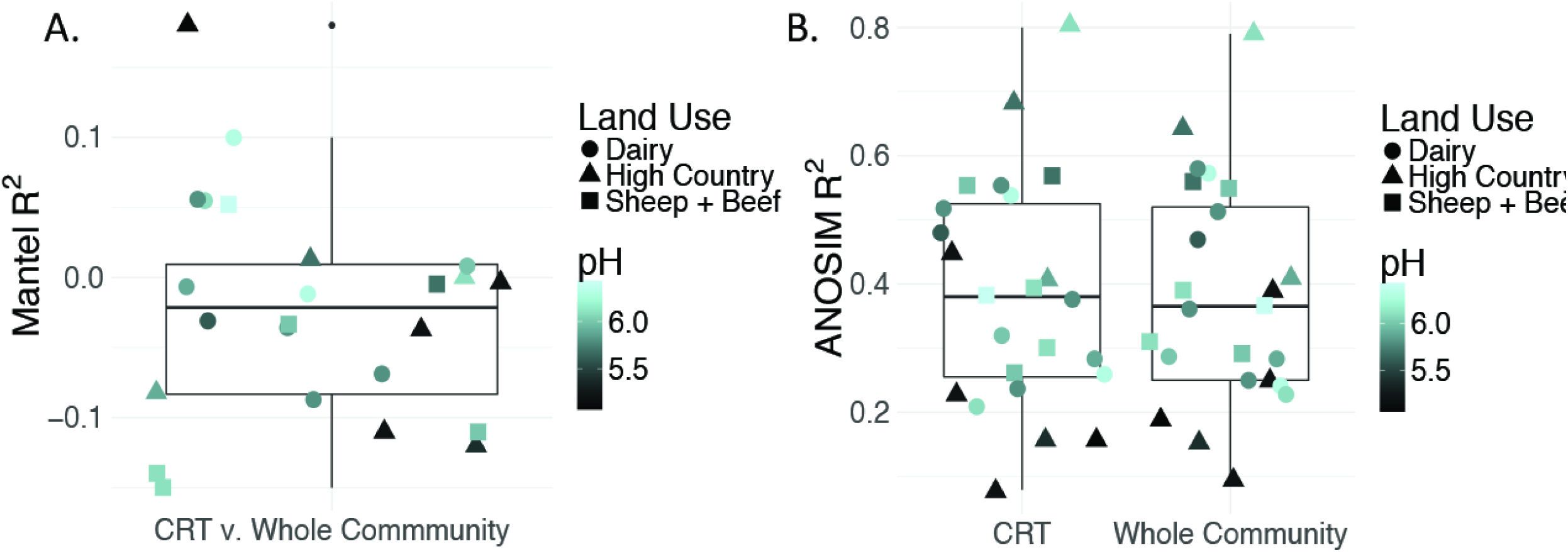
Contribution of CRT to community variance. Summary of Mantel correlations between Bray Curtis distances for site-level time responsive CRT communities and whole communities for each site (A) and summary of ANOSIM results for correlation between time and either site-level CRT or site-level whole community changes (B). pH and land use are shown to discount confounding effects by a dominant soil driver.

Despite a minor contribution to whole community variance, CRT community structure is linked to spatiotemporal factors. ANOSIM tests revealed significant correlations for CRT and whole communities at individual sites with time (Figure 1B, Figure S4-5). Mantel tests between space responsive CRT, the whole community and pH were significant, and ANOSIM tests with land use and soil order were also significant (Figure S6-7, Table S2). These results agree with previous studies which have found that soil prokaryotic communities exhibit temporal patterns (Lauber et al., 2013), are sensitive to pH change (Lauber et al., 2009), land use change (Steenwerth et al., 2003) and soil type (Kaminsky et al., 2017). This indicates that soil CRT follow the same assembly rules as abundant taxa, thus contributing to community changes, but are not overwhelmingly represented in these dynamics.

Although CRT communities exhibit broad relationships with spatiotemporal factors, results revealed that only 8% of time responsive CRT and 0.005% of space responsive CRT are correlated to measured spatiotemporal factors (Figure 2). Key taxa are represented; for example, *Acidobacteria* is widely known to be sensitive to pH, and is reflected as such here, where it is rare in high pH soils and abundant in low pH soils. *Saprospiraceae* vary seasonally, which is consistent with previous findings (Schauer et al., 2006). These results may show that certain rare members have major functional roles in soils, but aren’t well represented in overall community variance. Further, it shows that ecological filters not accounted for here, such as neutral processes or unmeasured niche factors, govern most soil CRT.

**Figure 2.**
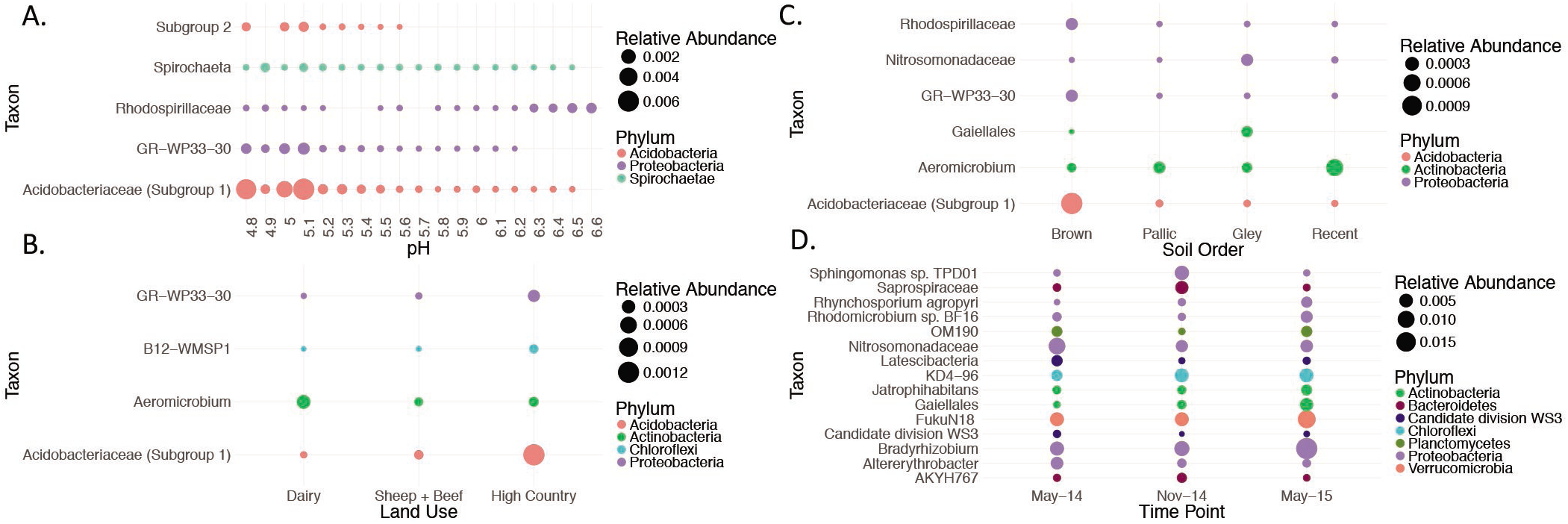
Relationships between individual CRT and key spatiotemporal factors. OTUs identified as CRT and significantly correlated to changes in pH (A), land use (B), soil order (C) and time (D). The OTU from each site that varied most significantly with time was plotted. A full list of OTUs correlated with time is reported in Table S3. Taxa were chosen based on a Kruskal Wallis *P* < 0.05 (land use, soil order and time), and a Spearman’s *P* < 0.05 for pH. Taxa are presented at the lowest classification level available, and colored at the phylum level.

Results indicate that while soil CRT are sensitive to spatiotemporal filters, they are not accountable for observed whole community variability across space and time. This is significant in that it implies an ecological role for soil CRT that is not related to abundance.

## Acknowledgements

We thank Mainland Minerals and Mainland Minerals Southern for sampling aid. We also thank Hill Laboratories and Soiltech for providing physicochemical analyses. RK was funded through a Callaghan Innovation education fellowship (MMSOU1301).

